# A supervised statistical learning approach for accurate *Legionella pneumophila* source attribution during outbreaks

**DOI:** 10.1101/133033

**Authors:** Andrew H. Buultjens, Kyra Y. L. Chua, Sarah L. Baines, Jason Kwong, Wei Gao, Zoe Cutcher, Stuart Adcock, Susan Ballard, Mark B. Schultz, Takehiro Tomita, Nela Subasinghe, Glen P. Carter, Sacha J. Pidot, Lucinda Franklin, Torsten Seemann, Anders Gonçalves Da Silva, Benjamin P. Howden, Timothy P. Stinear

## Abstract

Public health agencies are increasingly relying on genomics during Legionnaires’ disease investigations. However, the causative bacterium (*Legionella pneumophila*) has an unusual population structure with extreme temporal and spatial genome sequence conservation. Furthermore, Legionnaires’ disease outbreaks can be caused by multiple *L. pneumophila* genotypes in a single source. These factors can confound cluster identification using standard phylogenomic methods. Here, we show that a statistical learning approach based on

*L. pneumophila* core genome single nucleotide polymorphism (SNP) comparisons eliminates ambiguity for defining outbreak clusters and accurately predicts exposure sources for clinical cases. We illustrate the performance of our method by genome comparisons of 234 *L. pneumophila* isolates obtained from patients and cooling towers in Melbourne, Australia between 1994 and 2014. This collection included one of the largest reported Legionnaires’ disease outbreaks, involving 125 cases at an aquarium. Using only sequence data from *L. pneumophila* cooling tower isolates and including all core genome variation, we built a multivariate model using discriminant analysis of principal components (DAPC) to find cooling tower-specific genomic signatures, and then used it to predict the origin of clinical isolates. Model assignments were 93% congruent with epidemiological data, including the aquarium Legionnaires’ outbreak and three other unrelated outbreak investigations. We applied the same approach to a recently described investigation of Legionnaires’ disease within a UK hospital and observed model predictive ability of 86%. We have developed a promising means to breach *L. pneumophila* genetic diversity extremes and provide objective source attribution data for outbreak investigations.

## Importance

Microbial outbreak investigations are moving to a paradigm where whole genome sequencing and phylogenetic trees are used to support epidemiological investigations. It’s critical that outbreak source predictions are accurate, particularly for pathogens like *Legionella pneumophila*, which can spread widely and rapidly via cooling system aerosols causing Legionnaires’ disease. Here, by studying hundreds of *Legionella pneumophila* genomes collected over 21 years around a major Australian city, we uncovered limitations with the phylogenetic approach that could lead to misidentification of outbreak sources. We implement instead a statistical learning technique that eliminates the ambiguity of inferring disease transmission from phylogenies. Our approach takes geolocation information and core genome variation from environmental *L. pneumophila* isolates to build statistical models that predict with high confidence the environmental source of clinical *L. pneumophila* during disease outbreaks. We show the versatility of the technique by applying it to unrelated Legionnaires’ disease outbreaks in Australia and the UK.

## Introduction

Legionellae are Gram-negative bacteria that replicate within free-living aquatic amoebae and are present in aquatic environments worldwide. These bacteria can proliferate in man-made water systems and cause large outbreaks of pneumonia known as Legionnaires’ disease when contaminated water is aerosolized and inhaled (1). The majority of human infections are caused by *Legionella pneumophila* serogroup 1 (2). Public health investigations of Legionnaires’ disease outbreaks are typically supported by molecular typing methods to establish the likely source of the bacteria and the extent of the outbreak. Investigations usually proceed with the assumption that a single *Legionella* genotype is responsible for an environmental point source reservoir (3). Traditional molecular typing methods described for fingerprinting Legionellae include pulsed-field gel electrophoresis (PFGE) and sequence-based typing (SBT) (4). Increasingly, whole genome sequencing (WGS) is being employed to investigate individual *Legionella* outbreaks and the insights obtained from these high-resolution comparisons are challenging our expectations regarding common-source outbreaks, which usually are characterized by a single strain or genotype (5-9). It is becoming evident that outbreaks can be caused by multiple co-circulating *L. pneumophila* genotypes (5, 10) and that *L. pneumophila* core genomes can be surprisingly conserved across space and time (8, 11-13).

Melbourne is in the state of Victoria and it is the second largest city in Australia with a population approaching five million inhabitants, and considered the ninth largest city in the Southern Hemisphere. Legionellosis has been a notifiable disease in Victoria since 1979 and there are 50-100 cases reported each year, most occurring in the greater metropolitan region of Melbourne (14). The Microbiological Diagnostic Unit Public Health Laboratory (MDU PHL) is Victoria’s State Reference Laboratory for the characterization and typing of *Legionella* spp. The laboratory’s collection includes isolates from a particularly noteworthy outbreak at the Melbourne Aquarium in April 2000. This was the largest single episode of Legionellosis reported in Australia (15), approximately three months after the aquarium was opened to visitors, with construction of the site completed in December 1999. It resulted in 125 confirmed cases, with positive cultures obtained from 11 patients. Our isolate collection also spanned 28 other potential legionellosis outbreaks or infection clusters, for which at least one culture isolate had been obtained.

In this study, we used comparative genomics to explore the population structure of 234 *Legionella pneumophila* isolates recovered from human and environmental sources submitted to the MDU PHL in Melbourne over a 21-year period. This collection included 11 clinical and 14 environmental isolates from the Aquarium outbreak and 42 clinical and 50 environmental isolates from 28 other likely point source case clusters. We also assessed genomic data from a recently described investigation of Legionnaires’ cases at a UK hospital (8). The aim of this project was to develop a robust genomic approach that would surmount the unusual population structure of

*L. pneumophila* and assist identification of case clusters and source tracking efforts during Legionnaires’ disease outbreak investigations.

## Results

### Isolates and epidemiology

There were 234 *Legionella pneumophila* serogroup 1 (Lpn-SG 1) isolates obtained across a 21-year period between 1994 and 2014. Initial MLST analysis indicated that 180 isolates (77%) belonged to ST30. The collection comprised 180 clinical isolates of respiratory origin (sputum or bronchoscopy specimens) and 64 environmental isolates recovered from cooling tower water samples. All isolates were collected in the state of Victoria with the exception of six isolates from patients who were exposed elsewhere. Further information for each isolate is available in Table S1, including NCBI SRA accession numbers. One hundred and ten of the 234 isolates were epidemiologically associated with 29 formally investigated case clusters or outbreaks, designated as outbreaks A-AC (Table S1). The majority of these cases occurred within a 42 km radius of Melbourne city center and over a 16-year period. Outbreak A, the Melbourne Aquarium outbreak, was the largest (15).

### Complete genome sequence of *Legionella pneumophila* serogroup 1 isolate Lpm7613

Before this study, there were no closed, fully assembled ST30 *L. pneumophila* genomes. Thus, to ensure identification of maximum genetic variation among this dominant ST in our collection, we first established a ST30 reference genome sequence, selecting a clinical isolate from the Melbourne Aquarium outbreak (Lpm7613). The finished genome consisted of a single circular 3,261,562 bp chromosome (38.3% GC) and a 129,875 bp circular plasmid (pLpm7613) (Fig. S1). Although the chromosome indicated this genome belonged to the same lineage as *L. pneumophila* Philadelphia (Fig. 1A), the plasmid shared 100% nucleotide identity with pLPP reported in *L. pneumophila* Paris, but 2kb shorter in length (16). A total of 2,891 chromosomal protein-coding sequences (CDS), 43 tRNA genes and nine rRNA loci were predicted using Prokka (17). CRISPR-Cas regions were not detected (18).

**Fig. 1:**
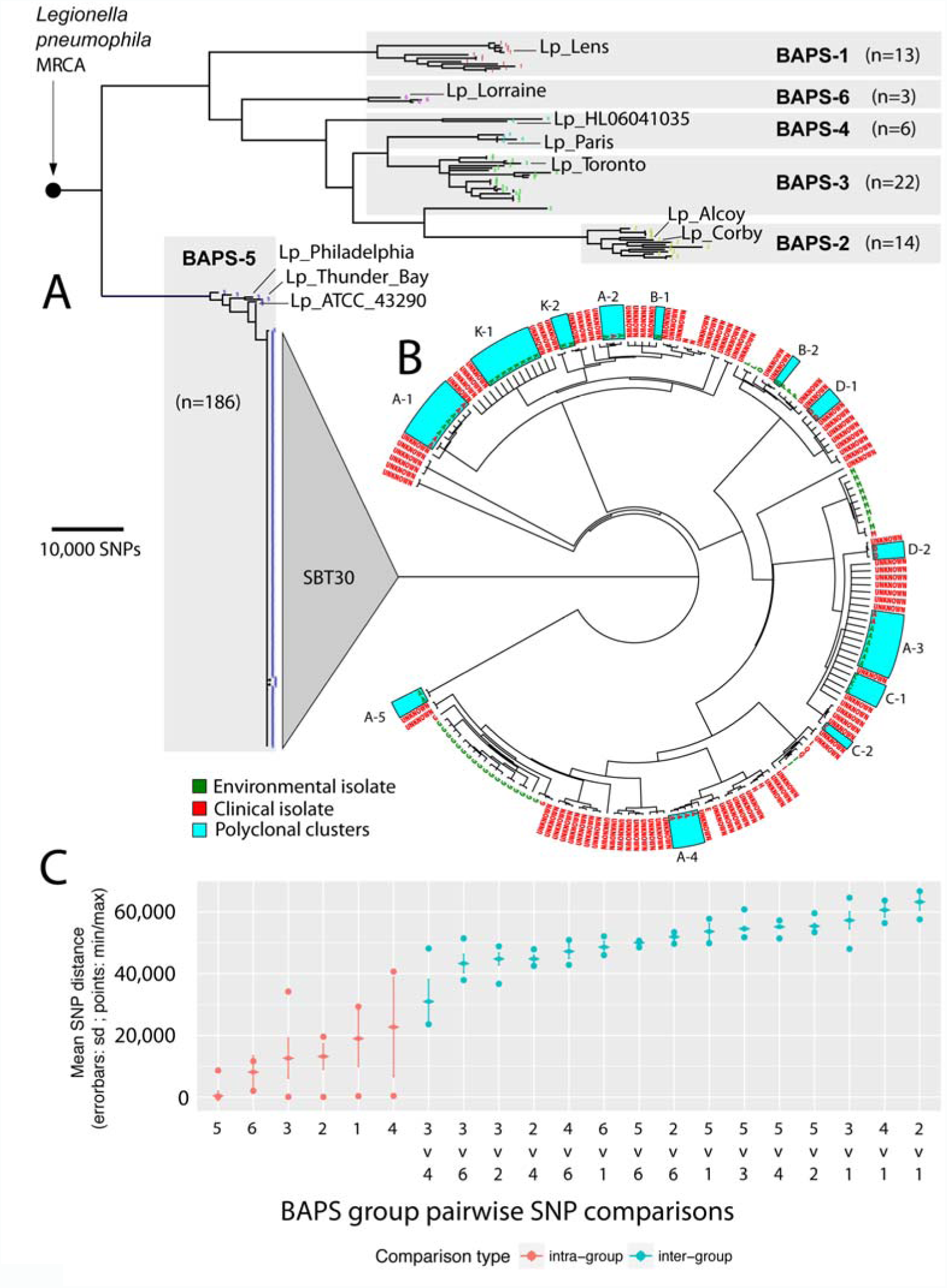
Global *Legionella pneumophila* population clustering, phylogenomics and genomic molecular epidemiology of local outbreaks. (A) Core genome phylogeny estimated using maximum likelihood corresponds with six BAPS groups. Branches with less than 70% bootstrap support were collapsed and scale indicates the number of core SNPs. The locations of the ten international genomes are labeled. (B) ST30 core genome phylogeny. Tree tips are labeled with outbreak codes. Environmental and clinical isolates are colored according to the key. Polyclonal outbreaks/case clusters are highlighted with blue boxes. Branch lengths have been transformed and are proportional to the number of nodes under each parent node. (C) Core genome pairwise SNP comparisons of within and between BAPS groups. All groups had smaller within group diversity compared to comparisons between groups.

### Assessment of L. pneumophila population structure

Sequence reads from the other 233 genomes and the ten selected publicly available completed genomes were mapped to the chromosome of reference strain Lpm7613. Approximately 90% of the Lpm7613 genome was present in all genomes (*i.e*., core), with 188,049 variable core nucleotide positions identified. Population structure analyses using an unsupervised Bayesian clustering approach revealed six distinct groups (BAPS groups) (Fig. 1A). Comparison of intra- and inter-BAPS group pairwise SNP distances confirmed the validity of these clusters and highlighted the extensive genetic variation among this Lpn-SG1 population (Fig. 1C). The exceptions were BAPS groups 3 and 4, which classified isolates across two clades, and is likely explained by recombination. Most striking, however, was the lack of diversity within the 186 genomes comprising BAPS group 5 (hereafter referred to as BAPS-5), with a median core SNP distance of only 5 SNPs (IQR 3 – 7). Isolates dispersed across time and space (including isolates from England, New South Wales, South Australia and Tasmania) were scattered throughout the entire phylogeny. All 180 ST30 isolates were encompassed by BAPS-5, as was ST37 *L. pneumophila* Philadelphia (Philadelphia, USA), ST211 *L. pneumophila* ATCC 43290 (Denver, USA) and ST733 *L. pneumophila* Thunder Bay (Ontario, Canada) (Fig. 1A,B). The median inter-BAPS group distances ranged between 27,506 to 63,136 SNPs (Fig. 1C), highlighting that there is also substantial genetic diversity among Lpn-SG1 isolates circulating in Melbourne.

A rooted maximum likelihood phylogeny of the population was then inferred using the 181,633 non-recombining core SNP loci. The phylogenomic tree reflected the BAPS clusters with BAPS-5 forming a distinct, well-supported lineage (Fig. 1A). The separation of the three North American reference isolates from the Melbourne ST30 isolates is suggestive of contemporaneous global dispersal of this BAPS-5 lineage (Fig. 1A,B). All BAPS groups displayed monophyletic origins with the exception of BAPS-3 and BAPS-4. BAPS-3 had a single isolate of paraphyletic origin that shared a most recent common ancestor (MRCA) with BAPS-2 while BAPS-4 contained two paraphyletic sub-clades, one of which shared a MRCA with the majority of BAPS-3 isolates.

### Impact of recombination

Recombination is a driving force in the evolution of the Legionellae (5, 7, 19-22). Therefore, to further understand the structure and evolution of this Lpn-SG1 population we assessed the impact of DNA exchange. There was evidence of extensive recombination among isolates across BAPS groups 1-4, and 6 with approximately 3% of variable nucleotide sites impacted relative to the Lpm7613 reference chromosome. The detection of two paraphyletic groups (BAPS-3 and BAPS-4) is likely explained by ancestral recombination among the component sub-clades. In comparison, there was little recombination evident among BAPS-5 isolates (Fig. S2), in accord with the core SNP phylogeny described above, and suggesting the relatively recent emergence of this *L. pneumophila* lineage. After removal of putative sequences affected by recombination, tree branch lengths showed no correlation with isolation dates (r^2^ =0.116). This observation indicates that nucleotide substitutions in the population have not been evolving under a molecular clock model, thus limiting estimates for dates of emergence for particular lineages.

### Genomic molecular epidemiology of local outbreaks

We next compared only the 180 ST30 genomes to our Lpm7613 reference genome, and again confirmed the very restricted genomic diversity within this lineage (median core SNP distance was 6 SNPs (IQR 4 – 9), with five outlier genomes, impacted by recombination. (Fig. 1B,C Fig. S2). Within this reconstructed core-genome ST30-specific phylogeny, many but not all epidemiologically-related isolates formed distinct, well-supported, monophyletic clades. In some instances, epidemiologically-associated isolates spanned multiple clades (outbreaks A, B, C, D and K) (Fig. 1B). In addition, Outbreak A (the Melbourne aquarium outbreak), which was previously considered to represent infections caused by a single clone (Table S1) (15), actually contained five distinct genotypes (A1-A5) (Fig 1B,C).

The analysis of environmental surveillance isolates provided an ideal means to gain insights into the diversity within potential reservoirs of Lpn-SG1 - diversity that might enable prospective source tracking. A phylogeographic analysis was therefore undertaken to assess the relationship between 64 environmental Melbourne metropolitan isolates against their 11 cooling tower sampling locations. Based on variation in core SNPs, striking geographical structure was observed, with the majority of isolates from common cooling towers tightly clustering in the phylogeny (Fig. 2A). Comparisons of pairwise core SNPs depicted smaller within group diversity and larger between location group diversity, further indicating the existence of geographical population structure (Fig. 2B). This structure among the environmental Lpn-SG1 isolates suggested it might be possible to use the genome data to build models predictive of environmental source to assist epidemiological efforts during outbreak investigations.

**Fig. 2:**
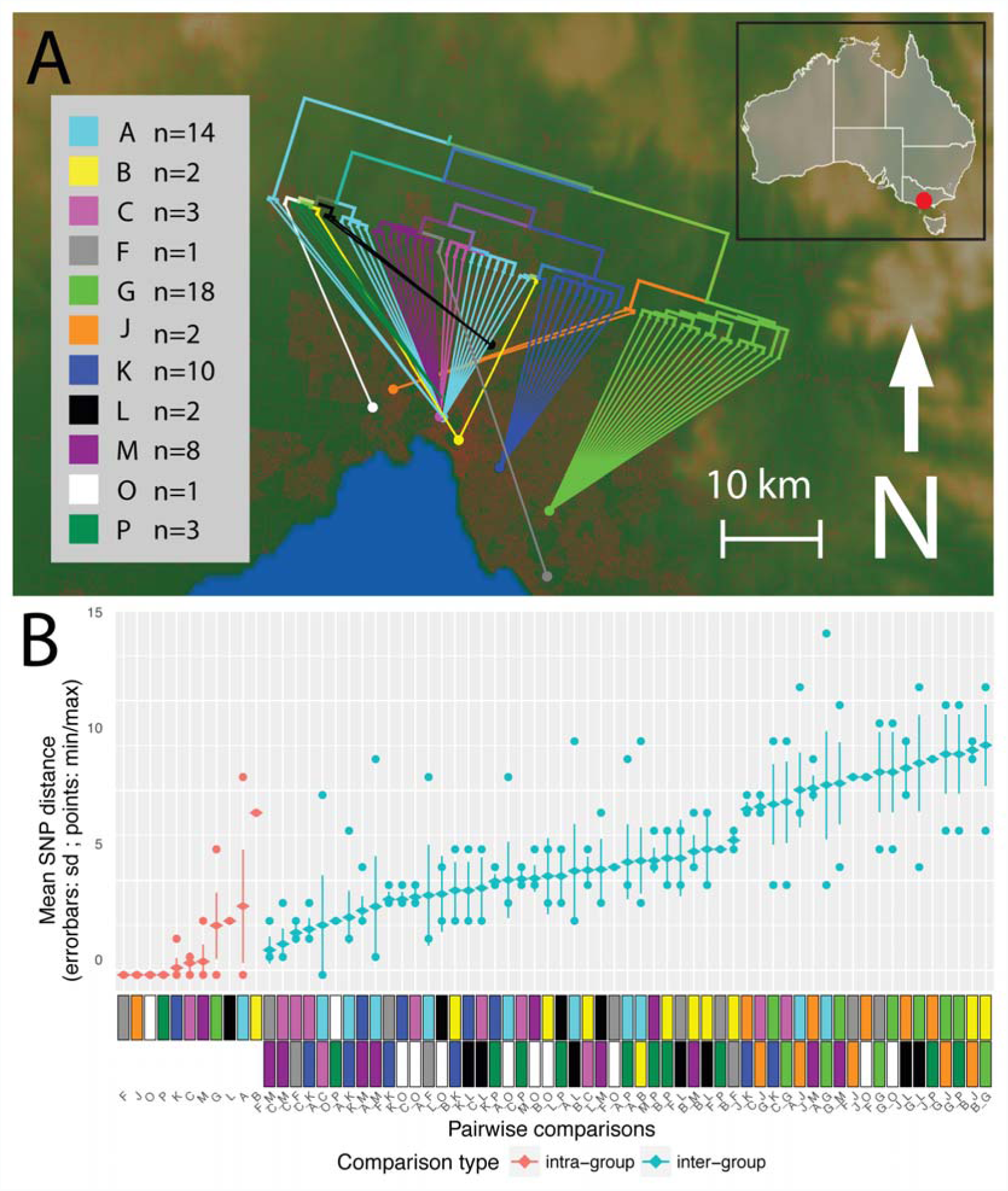
Phylogeography of 64 Lpn-SG1 environmental isolate genomes. (A) Map of the greater Melbourne area, showing the location of the 11 cooling towers assessed during Legionellosis outbreaks, designated by colored circles. ‘A’ (light blue) represents the location Melbourne aquarium outbreak and is close to the centre of Melbourne. Inset shows the location of Melbourne (red circle) within the State of Victoria in south east Australia. Overall the phylogeny aligns closely with the geography of originating cooling towers. For several outbreak codes polyclonality is apparent, as some common origins have connecting lines drawn from different sub-clades of the phylogeny. Red coloration on the base map represents population density within the greater Melbourne region. The branch lengths of the trees have been transformed and are proportional to the number of nodes under each parent node. (B) Core genome pairwise SNP comparisons of within, and between, cooling tower isolate groups. Comparisons of specific epidemiologically defined groups (infection clusters) are indicated with color codes as defined in the key. All groups had smaller within diversity than between group diversity.

### A multivariate statistical model for source attribution

To enhance resolution and try to detect outbreak-specific genomic signals, a supervised statistical learning approach called Discriminant Analysis of Principal Components (DAPC) (23) was employed. DAPC is a linear discriminant analysis (LDA) that accommodates discrete genetic-based predictors by first transforming the genetic data into continuous Principal Components (PC) and building predictive classification models. The PCs are used to build discriminant functions (DF) under the constraint that they must minimize within group variance, and maximize variance between groups. Infection clusters were defined *a priori* from the epidemiological findings, and training (environmental) isolates were used to establish the discriminant functions. The model was then be used to estimate the posterior probability of membership for an unknown (*e.g.* clinical) isolate for each pre-specified infection cluster given the training data. Here, we used 43 of the 64 environmental isolates in the training set (cooling tower isolates originating from epidemiologically defined infection clusters that possessed at least one environmental and clinical representative), under the assumption that each outbreak was caused by exposure to a point source of Lpn-SG1. We used core genome SNPs from only environmental Lpn-SG1 genomes to build the classifier (24).

Outbreak-associated environmental Lpn-SG1 were grouped *a priori* into training set groups based on the origin of the cooling towers from which they were isolated (see model building details in methods). The DFs were then used to classify 15 clinical isolates that had been independently assigned based on epidemiological data to the training set groups, hereon referred to as the validation genomes (Table S1). The input matrix for DAPC was an alignment of 714 non-recombinogenic SNPs variable among the 43 environmental genomes. Plots depicting the separation of isolates according to the first two discriminant functions and the amount of variation explained is shown (Fig. 3). A model was trained using the first four principal components (PC), as this was found to be optimal (see methods). We next classified our clinical validation genomes using the model and found a 93% match between our model’s assignment and that proposed by the epidemiological data (Fig. 4A,B). These data show that despite the high level of genome conservation and the presence of multiple genotypes within a single environmental source, it is possible to utilize signature differences in core genome SNPs to build predictive probabilistic classification models. The single discrepancy between model predictions and epidemiological groupings was an infection cluster C genome that was predicted as originating from the Melbourne Aquarium. Interestingly, cluster C was located closest to the Melbourne Aquarium at a distance of approximately 500 metres. Given the proximity of clusters A and C, these data may indicate cooling towers were seeded from a common *L. pneumophila* source. In order to appraise the utility of this method beyond a large urban setting and the ST30 genotype, we built a sister model using 31 ST1 environmental *L. pneumophila* genomes from a previously published hospital investigation in Essex, UK, and used it to predict the origins of seven nosocomial clinical isolates (Fig. 3A, Table S1) (8). Here, the model was trained using an alignment of 59 non-recombinogenic SNPs among the 31 environmental genomes and retaining the first 15 PCs, as this was found to be optimal. As with the Melbourne disease clusters, the model performed very well. For 86% of the clinical isolates there was a match between the model’s ward assignment and the origin suggested by epidemiology (Fig. 4A,B). Again, a single discrepancy occurred with a ward G genome predicted to originate from ward A. Wards A and G were co-located on the same corner and level of a common building, again suggesting a common *L. pneumophila* source (8). As before, isolates from a common source would be miss-assigned by the model, owing to the lack of location-specific genomic variants.

**Fig. 3.**
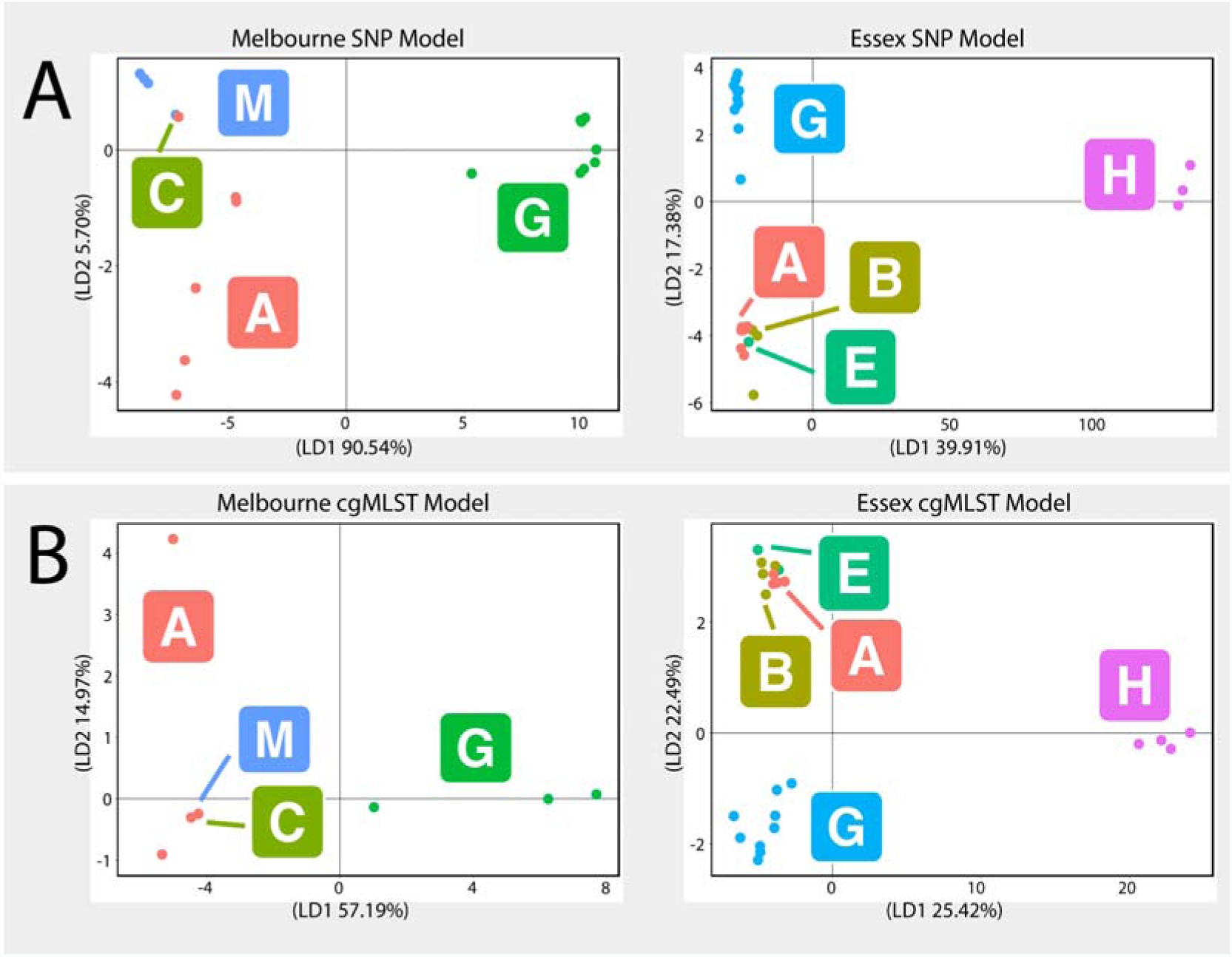
Scatterplots resulting from discriminant analysis of principal components (DAPC). (A): Core genome single nucleotide polymorphisms (SNP) based models of the Melbourne (top left) and Essex (top right) datasets and; (B): core genome MLST (cgMLST) based models of the Melbourne (bottom left) and Essex (bottom right) datasets. The membership of each point within an epidemiologically defined cluster (*e.g.* “A” is the Melbourne Aquarium outbreak) is indicated by the colored circles and the corresponding letters labeled within squares. The amount of variation explained by the first and second discriminant functions are specified on the axes of each plot.

### Core genome multilocus sequence typing (cgMLST) has reduced discrimination

In order to evaluate the utility of the recently described cgMLST scheme for source tracking (25, 26), we trained new DAPC models for both the Melbourne and Essex hospital datasets using a matrix of allelic integers derived from SNP profiles of the 1,529 cgMLST loci (Fig. 3B). When using the first one and seven PCs, we observed only 60% and 71% concordance between our model’s assignment and that predicted by the epidemiological data for the Melbourne and Essex hospital datasets, respectively (Fig. 4A,C).

**Fig. 4:**
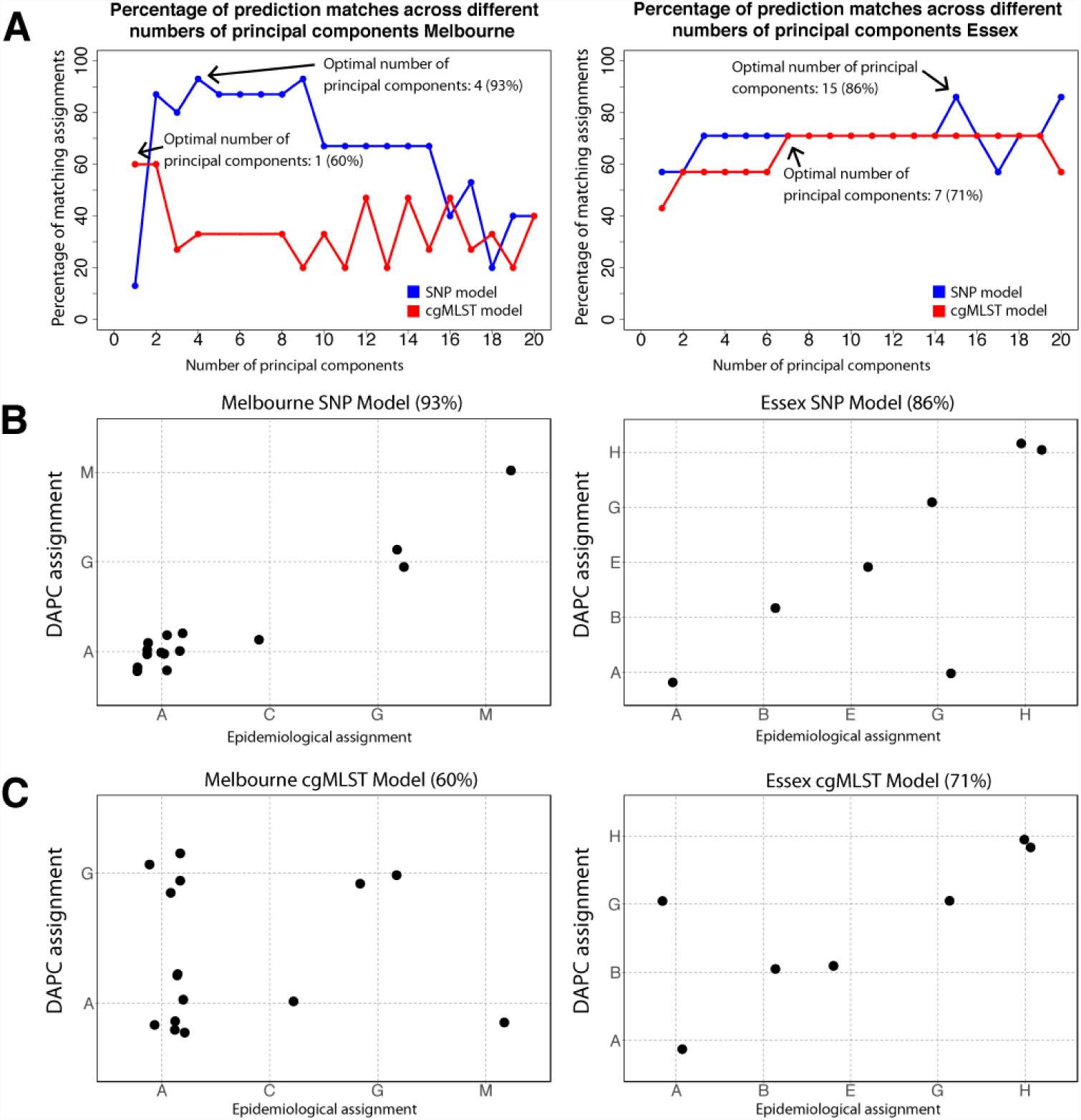
Discriminatory analysis of principal components (DAPC) modeling of Lpn-SG1 genomic data. (A) Model comparison plots depicting the percentage of matches between the predicted and epidemiologically determined groupings of the validation set genomes across a range of 1-20 principal components for single nucleotide polymorphisms (SNP) and core genome MLST (cgMLST) DAPC models for the Melbourne and Essex datasets. The retention of four and one principal components was found to be optimal for the SNP (93% match) and cgMLST (60% match) models in Melbourne, respectively, while 15 and seven principal components were found to be optimal for the SNP (86% match) and cgMLST (71% match) models in the Essex hospital, respectively. (B) Assignment plots depicting the ability of the SNP models to predict the source attribution of the validation set clinical isolate genomes for the Melbourne and Essex hospital datasets. (C) Assignment plots depicting the ability of the cgMLST models to predict the source attribution of the validation set clinical isolate genomes for the Melbourne and Essex hospital datasets.

## Discussion

In this study, we have retrospectively examined a large collection of 234 clinical and environmental isolates Lpn-SG1 isolates spanning 29 defined outbreaks. Isolates were collected over wide temporal and spatial scales and detailed genomic comparisons revealed wide extremes Lpn-SG1 genetic diversity among distinct genomic populations; a phenomenon not fully appreciated from previous genomic investigations that have sampled less extensively and focused on single outbreaks (5, 6, 27). Most striking in our collection was the high sequence conservation and dominance of a single genotype (BAPS-5, ST30), shared by 77% of isolates with a median core SNP distance of only 5 SNPs across 21 years. In agreement with our findings, two recent population genomic investigations of Lpn-SG1 also describe the unusual restriction in core genome diversity (8, 12).

Based on our previous experience with other bacterial pathogens (28) and reports in the literature of Legionnaires’ disease outbreak investigations using genomics (27, 29) - we expected to be able to use Lpn-SG1 genomic comparisons and develop genetic rule-in or rule-out criteria to guide outbreak assessment and source attribution. For example, we recently proposed a ‘traffic-light’ system for *Listeria monocytogenes* based on SNP difference cutoffs of ‘likely related’, ‘possibly-related’ and ‘not-related’ (28). This approach has also been proposed for *L. pneumophila* (25). A comparison of genotyping approaches using 335 Lpn isolates, including 106 from the European Society for Clinical Microbiology Study Group’s *Legionella Typing Panel*, proposed an escalating, hierarchical approach to genotyping, beginning with an extended 50-gene MLST scheme up to a 1529-gene cgMLST (25, 26).

The analysis of the population structure of Lpn-SG1 presented here indicates that SNP-based typing with threshold cut-offs, whether they are based on seven genes, 50 genes, 1500 genes or whole genomes will not necessarily provide sufficient discriminative power. These genotyping approaches will be confounded by the presence of (i) indistinguishable Lpn-SG1 genotypes present in unrelated cases and (ii) polyclonal outbreaks. Our retrospective analysis of the Melbourne Aquarium outbreak illustrates clearly both these issues, where five distinct subtypes were recovered from 25 clinical and environmental isolates (Fig. 1B). There is a growing awareness of single source, polyphyletic Lpn-SG1 outbreaks (8, 10, 13, 30, 31). These data all point to the need for a different approach in order to use molecular epidemiology and genomics in support of *Legionella* outbreak investigations.

We address this issue by exploiting all core genome information to train probabilistic classification models. Our DAPC analysis demonstrates that it is possible to build predictive models based on Lpn-SG1 environmentally derived genomes that help in identifying the source of clinical isolates during complex outbreak investigations in both the community and hospital environments (Fig. 4). By including all core SNP variation, DAPC was able to identify outbreak-specific genotypes, even when the source of the outbreak was polyclonal. This enabled us to build robust models that assigned validation set genomes, with known provenance, back to their original groupings with high concordance. The fact that this model was built purely from environmental surveillance isolates demonstrates that such approaches can be developed prospectively and be preexisting, ready to deploy at the onset of outbreaks.

In contrast to the high performance of the DAPC model developed from core genome SNPs, the model built using variants identified by cgMLST scheme had a lower matching rate when assigning validation genomes back to their putative epidemiological groupings (Fig. 4C). Despite cgMLST being a useful tool for broad Lpn-SG1 population structure assessment, our analysis suggests it may have insufficient resolution and thus predictive capacity for outbreak investigations.

The DAPC approach however, while promising, does not permit discrimination among isolates that do not belong to defined clusters. This is because the model assumes that the world is composed of only the *k* groups used to train it, and therefore assigns unknown isolates to one of these groups, even if the isolate is known not to be part of any of the groups. One way to address this issue would be to create a single group classifier, which is trained with environmental samples. Isolates with low probability of membership to this single large group would then be excluded before being analyzed with the multi-group model. Future models could be further improved by adding epidemiological evidence (*e.g.* patient zip codes), and assess how that improves our assignment of a clinical isolate to a particular location. An advantage of a classification-based model is that its output could be distilled down to a zip code (or group of zip codes) and a probability that a clinical isolate is associated with the zip code (indicating uncertainty about the classification). This would obviate the need to interpret, and explain phylogenetic trees. Interpreting trees is often not intuitive and trees may fail to communicate what action is required from a public health perspective. Crucial for such a classification approach to work however, is an extensive and temporally dynamic database of environmental Lpn-SG1 genotypes. That is, there would need to be ongoing surveillance and isolation of Lpn-SG1 from environmental sources. We are currently investigating how to implement such models.

The modeling approach, is not intended to be used in isolation, but rather employed as an adjunct to traditional epidemiological investigations. In this way, insights gained through epidemiological investigations can be informed by microbiological evidence from our predictive models. A limitation of our current models are the relatively small sample sizes. Performance measures for validation sets this small are often sensitive to slight perturbations in the data and may be influenced by small features of the data. However, as a proof-of-concept implementation of our approach, we have built two models from independent datasets, and both demonstrate high predictive capacity. More robust appraisals of model performance will require validation with larger datasets, collected prospectively.

From a biological perspective, the lack of genetic diversity in Lpn-SG1 over such coarse temporal and spatial scales is potentially explained by a reservoir of latent-state bacteria intermittently seeding warm water sources in the greater Melbourne region and is supported by the frequently-reported and widespread presence of *Legionella* species in drinking water supply systems (DWSS) (32-34). Independent studies propose similar hypotheses to explain the surprisingly high sequence conservation among some *L. pneumophila* genomes (8, 12).

This study is, to our knowledge, the largest genomic investigation of environmental and clinical Legionella reported to date from a single jurisdiction and confirms that Lpn-SG1 is an unusual ‘edge case’ in the application of genomics in public health microbiology. In the absence of a deep understanding of local *L. pneumophila* population structure (both clinical and environmental) the combination of extreme genomic monomorphism combined with outbreaks caused by mixed pathogen populations could easily lead to erroneous conclusions regarding source attribution. Thus, we require new approaches that can better utilize the genomic information available, and harmoniously combine it with epidemiological evidence, in order to provide public health officials with useful and timely information.

## Materials and Methods

### Bacterial strains, growth conditions, case definitions

*Legionella pneumophila* serogroup 1 isolates were resuscitated from -80°C storage and assessed. Duplicate isolates from the same patient were excluded from the study. Isolates were cultured for 48-72 h at 37°C on BCYE agar and re-confirmed serogroup 1 by latex agglutination (Oxoid). Metadata collected on all isolates included year of isolation and country or city of isolation. Cases resident in the state of Victoria, Australia, were assessed by the Victorian State Government public health unit in accordance with national guidelines and an outbreak investigation was initiated when common exposures were reported by different cases whose onset dates occurred within a two-week window. (http://www.health.gov.au/internet/main/publishing.nsf/content/cdna-song-legionella.htm, accessed 31 August 2015). In this manner, we were able to determine the human cases epidemiologically linked to each other. Many of the outbreaks/infection clusters contained a greater number of cases than there were isolates as the diagnosis of Legionellosis was made by culture-independent methods. Complete, closed genomes of *L. pneumophila* that were publicly available were obtained from GenBank for inclusion in the analysis (Table S1).

### Sequence based typing

This was performed as previously described according to the European Legionnaires’ Disease Surveillance Network (ELDSNet) method (http://bioinformatics.phe.org.uk/legionella/legionella_sbt/php/sbt_homepage.php, accessed 31 August 2015) (35).

### DNA sequencing

DNA libraries were prepared using the NexteraXT DNA preparation kit (Illumina) and whole genome sequencing was performed on the NextSeq platform (Illumina) with 2x150 bp chemistry. For single molecule real-time (SMRT) sequencing (Pacific Biosciences), genomic DNA was extracted from agarose plugs using the CDC Pulsenet Protocol to allow for recovery of high molecular weight, intact DNA (http://www.cdc.gov/pulsenet/pathogens, accessed 31 August 2015). Size-selected 10kb DNA libraries were prepared according to manufacturers’ instructions and sequenced on the RS II platform (Pacific Biosciences) using P6-C4 chemistry. All sequence reads and the completed genome are available (GenBank BioProject ID: PRJEB13594)

### *Legionella pneumophila* serogroup 1 isolate Lpm7613 assembly and closure

A high quality finished ST 30 reference genome was established for *L. pneumophila* serogroup 1 clinical isolate Lpm7613 using the SMRT(®) Analysis System v2.3.0.140936 (Pacific Biosciences). Raw sequence data were *de novo* assembled using the HGAP v3 protocol with a genome size of 4 Mb. Polished contigs were error corrected using Quiver v1. The resulting assembly was then checked using BridgeMapper v1 in the SMRT(®) Analysis System, and the consensus sequence corrected with short-read Illumina data, using the program Snippy (https://github.com/tseemann/snippy). Whole genome annotation was performed using Prokka (17), preferentially using the *L. pneumophila* Paris strain annotation (16). BRIG was used to visualize BLASTn DNA:DNA comparisons of *L. pneumophila* Lpm7613 against other *L. pneumophila* genomes (36). Nomenclature of the genomic islands demonstrated in *L. pneumophila* Lpm7613 was based on previously described islands (37). CRISPR databases were used to search for CRISPR sequences (http://crispi.genouest.org and http://crispr.u-psud.fr/Server/, accessed 14 February 2016).

### Variant detection and phylogenetic analysis

The genomes of ten publicly available complete *L. pneumophila* genomes (Table S1) were shredded to generate short *in silico* sequence reads of 250bp and all 244 *L. pneumophila* reads sets were mapped against the Lpm7613 reference genome using Snippy v3.2. An alignment file from pairwise comparisons of core genome SNPs (with inferred recombining sites removed) was used as input to FastTree v2.1.8 with double precision (38) to infer a maximum likelihood phylogenetic tree using the general time reversible model of nucleotide substitution. Branch support was estimated using 1,000 bootstrap replicates. Resulting trees were visualized in FigTree v1.4.2 (http://tree.bio.ed.ac.uk/software/figtree/). Single nucleotide polymorphism (SNP) differences between isolates were tabulated and visualized using a custom R-script (https://github.com/MDU-PHL/pairwise_snp_differences). The core genome SNPs were also used as the input into a Bayesian analysis of population structure (BAPS) using iterative clustering to a depth of 10 levels and a pre-specified maximum of 20 clusters (39).

### Recombination and molecular clock analysis

Recombination detection was performed using ClonalFrameML (40), taking as input a full genome alignment (included invariant sites) prepared using Snippy as above and the ML phylogeny as a guide tree with polytomies removed from the FastTree tree using a custom python script (https://github.com/kwongj/nw_multi2bifurcation). Results were visualized using a custom Python script to render separate and superposable images of extant and ancestral inferred recombination regions (https://github.com/kwongj/cfml-maskrc). Molecular clock-likeness of the ML tree with ClonalFrameML-adjusted branch lengths was assessed using TempEst v1.5 (http://tree.bio.ed.ac.uk/software/tempest/).

### Phylogeographic analysis

Variant detection for the 64 environmental genomes was undertaken by running snippy-core. Core SNPs were used to reconstruct a phylogenomic tree with FastTree that was overlaid upon a base map in GenGIS (41). Victorian population mesh data was downloaded from the Australia Bureau of Statistics webpage (http://www.abs.gov.au/AUSSTATS/abs@.nsf/DetailsPage/1270.0.55.001July%202016?OpenDocument) and Local Government Area data was downloaded from the Victorian Government Data Directory webpage (https://www.data.vic.gov.au/data/dataset/lga-geographical-profiles-2014-beta/resource/f6c49074-0679-4c79-a0db-04dac8eda364).

### DAPC model building using core SNPs

Discriminant analysis of principal components (DAPC) is a multivariate method that tries to reconstruct hypothesized subdivisions in a given population (typically formed from demographic or phenotypic information) using genomic data (42). DAPC was implemented in the R package *adegenet* v2.0.1 (42). For input, we used a matrix of single nucleotide polymorphisms (SNP) for all genomes originating from infection clusters that possessed at least one environmental and clinical representative (Table S1). SNP detection was undertaken by running Snippy and sites that were recombinogenic and or invariant among the environmental genomes were discarded. An input SNP matrix of exclusively environmental isolates (hereon referred to as the training set) was used to develop a DAPC model. The training set subdivisions were based on the geographic origin of the environmental isolates (Table S1) (23). The resultant model was then tested using clinical isolates (hereon referred to as the validation set). The ability of the model to predict the environmental source of the validation set genomes was simulated across the first to the 20th principal components, allowing an optimal number of principal components to be identified. The optimized model was then used to predict the environmental origin of the clinical isolate genomes.

### DAPC model building using cgMLST variation

In order to detect variants within the recently described cgMLST regions, reads were mapped to the Lp_Philadelphia chromosome (NC_002942.5) using snippy. SNP profiles from within the cgMLST regions were reduced to allelic integers, with all genes containing zero coverage or uncertain base-calls, excluded. Allelic integers were concatenated into a matrix and, using the same DAPC model-building method as mentioned above, models were established using the training set environmental genomes and used to predict the origin of the validation set clinical isolate genomes.

## References

1. Fields BS, Benson RF, Besser RE. 2002. Legionella and Legionnaires’ disease: 25 years of investigation. Clin Microbiol Rev 15:506–526.

2. Yu VL, Plouffe JF, Pastoris MC, Stout JE, Schousboe M, Widmer A, Summersgill J, File T, Heath CM, Paterson DL, Chereshsky A. 2002. Distribution of *Legionella species* and serogroups isolated by culture in patients with sporadic community-acquired legionellosis: an international collaborative survey. J Infect Dis 186:127–128.

3. Luck C, Fry NK, Helbig JH, Jarraud S, Harrison TG. 2013. Typing methods for Legionella. Methods Mol Biol 954:119–148.

4. Mercante JW, Winchell JM. 2015. Current and emerging Legionella diagnostics for laboratory and outbreak investigations. Clin Microbiol Rev 28:95–133.

5. McAdam PR, Vander Broek CW, Lindsay DS, Ward MJ, Hanson MF, Gillies M, Watson M, Stevens JM, Edwards GF, Fitzgerald JR. 2014. Gene flow in environmental Legionella pneumophila leads to genetic and pathogenic heterogeneity within a Legionnaires’ disease outbreak. Genome Biol 15:504.

6. Reuter S, Harrison TG, Koser CU, Ellington MJ, Smith GP, Parkhill J, Peacock SJ, Bentley SD, Torok ME. 2013. A pilot study of rapid whole-genome sequencing for the investigation of a Legionella outbreak. BMJ Open 3.

7. Sanchez-Buso L, Comas I, Jorques G, Gonzalez-Candelas F. 2014. Recombination drives genome evolution in outbreak-related Legionella pneumophila isolates. Nat Genet 46:1205–1211.

8. David S, Afshar B, Mentasti M, Ginevra C, Podglajen I, Harris SR, Chalker VJ, Jarraud S, Harrison TG, Parkhill J. 2017. Seeding and Establishment of Legionella pneumophila in Hospitals: Implications for Genomic Investigations of Nosocomial Legionnaires’ Disease. Clin Infect Dis 64:1251–1259.

9. Weiss D, Boyd C, Rakeman JL, Greene SK, Fitzhenry R, McProud T, Musser K, Huang L, Kornblum J, Nazarian EJ, Fine AD, Braunstein SL, Kass D, Landman K, Lapierre P, Hughes S, Tran A, Taylor J, Baker D, Jones L, Kornstein L, Liu B, Perez R, Lucero DE, Peterson E, Benowitz I, Lee KF, Ngai S, Stripling M, Varma JK, South Bronx Legionnaires’ Disease Investigation T. 2017. A Large Community Outbreak of Legionnaires’ Disease Associated With a Cooling Tower in New York City, 2015. Public Health Rep 132:241–250.

10. Sanchez-Buso L, Guiral S, Crespi S, Moya V, Camaro ML, Olmos MP, Adrian F, Morera V, Gonzalez-Moran F, Vanaclocha H, Gonzalez-Candelas F. 2015. Genomic Investigation of a Legionellosis Outbreak in a Persistently Colonized Hotel. Front Microbiol 6:1556.

11. Underwood AP, Jones G, Mentasti M, Fry NK, Harrison TG. 2013. Comparison of the Legionella pneumophila population structure as determined by sequence-based typing and whole genome sequencing. BMC Microbiol 13:302.

12. David S, Rusniok C, Mentasti M, Gomez-Valero L, Harris SR, Lechat P, Lees, J., Ginerva C, Glaser C, Ma L, Bouchier C, Underwood A, Jarraud S, Harrison TG, Parkhill J, Buchrieser C. 2016. Multiple major disease-associated clones of Legionella pneumophila have emerged recently and independently. Genome Res In press.

13. Schjorring S, Stegger M, Kjelso C, Lilje B, Bangsborg JM, Petersen RF, David S, Uldum SA, Infections ESGfL. 2017. Genomic investigation of a suspected outbreak of Legionella pneumophila ST82 reveals undetected heterogeneity by the present gold-standard methods, Denmark, July to November 2014. Euro Surveill 22.

14. Anon. 2014. Communicable Disease Surveillance, Victoria, Oct-Dec 2014. Victorian Infectious Diseases Bulletin 17:22–23.

15. Greig JE, Carnie JA, Tallis GF, Ryan NJ, Tan AG, Gordon IR, Zwolak B, Leydon JA, Guest CS, Hart WG. 2004. An outbreak of Legionnaires’ disease at the Melbourne Aquarium, April 2000: investigation and case–control studies. Med J Aust 180:566–572.

16. Cazalet C, Rusniok C, Bruggemann H, Zidane N, Magnier A, Ma L, Tichit M, Jarraud S, Bouchier C, Vandenesch F, Kunst F, Etienne J, Glaser P, Buchrieser C. 2004. Evidence in the Legionella pneumophila genome for exploitation of host cell functions and high genome plasticity. Nat Genet 36:1165–1173.

17. Seemann T. 2014. Prokka: rapid prokaryotic genome annotation. Bioinformatics 30:2068–2069.

18. Rao C, Guyard C, Pelaz C, Wasserscheid J, Bondy-Denomy J, Dewar K, Ensminger AW. 2016. Active and Adaptive Legionella CRISPR-Cas reveals a recurrent challenge to the pathogen. Cell Microbiol doi:10.1111/cmi.12586.

19. Coscolla M, Gonzalez-Candelas F. 2007. Population structure and recombination in environmental isolates of Legionella pneumophila. Environ Microbiol 9:643–656.

20. Coscolla M, Gonzalez-Candelas F. 2009. Comparison of clinical and environmental samples of Legionella pneumophila at the nucleotide sequence level. Infect Genet Evol 9:882–888.

21. Gomez-Valero L, Rusniok C, Jarraud S, Vacherie B, Rouy Z, Barbe V, Medigue C, Etienne J, Buchrieser C. 2011. Extensive recombination events and horizontal gene transfer shaped the Legionella pneumophila genomes. BMC Genomics 12:536.

22. Costa J, Teixeira PG, d’Avo AF, Junior CS, Verissimo A. 2014. Intragenic recombination has a critical role on the evolution of Legionella pneumophila virulence-related effector sidJ. PLoS One 9:e109840.

23. Jombart T, Devillard S, Balloux F. 2010. Discriminant analysis of principal components: a new method for the analysis of genetically structured populations. BMC Genet 11:94.

24. McNally A, Oren Y, Kelly D, Pascoe B, Dunn S, Sreecharan T, Vehkala M, Valimaki N, Prentice MB, Ashour A, Avram O, Pupko T, Dobrindt U, Literak I, Guenther S, Schaufler K, Wieler LH, Zhiyong Z, Sheppard SK, McInerney JO, Corander J. 2016. Combined Analysis of Variation in Core, Accessory and Regulatory Genome Regions Provides a Super-Resolution View into the Evolution of Bacterial Populations. PLoS Genet 12:e1006280.

25. David S, Mentasti M, Tewolde R, Aslett M, Harris SR, Afshar B, Underwood A, Fry NK, Parkhill J, Harrison TG. 2016. Evaluation of an optimal epidemiologic typing scheme for Legionella pneumophila with whole genome sequence data using validation guidelines. J Clin Microbiol doi:10.1128/JCM.00432-16.

26. Moran-Gilad J, Prior K, Yakunin E, Harrison TG, Underwood A, Lazarovitch T, Valinsky L, Luck C, Krux F, Agmon V, Grotto I, Harmsen D. 2015. Design and application of a core genome multilocus sequence typing scheme for investigation of Legionnaires’ disease incidents. Euro Surveill 20.

27. Bartley PB, Ben Zakour NL, Stanton-Cook M, Muguli R, Prado L, Garnys V, Taylor K, Barnett TC, Pinna G, Robson J, Paterson DL, Walker MJ, Schembri MA, Beatson SA. 2016. Hospital-wide Eradication of a Nosocomial Legionella pneumophila Serogroup 1 Outbreak. Clin Infect Dis 62:273–279.

28. Kwong JC, Mercoulia K, Tomita T, Easton M, Li HY, Bulach DM, Stinear TP, Seemann T, Howden BP. 2016. Prospective Whole-Genome Sequencing Enhances National Surveillance of Listeria monocytogenes. J Clin Microbiol 54:333–342.

29. Graham RM, Doyle CJ, Jennison AV. 2014. Real-time investigation of a Legionella pneumophila outbreak using whole genome sequencing. Epidemiol Infect 142:2347–2351.

30. Sanchez-Buso L, Olmos MP, Camaro ML, Adrian F, Calafat JM, Gonzalez-Candelas F. 2015. Phylogenetic analysis of environmental Legionella pneumophila isolates from an endemic area (Alcoy, Spain). Infect Genet Evol 30:45–54.

31. Coscolla M, Fernandez C, Colomina J, Sanchez-Buso L, Gonzalez-Candelas F. 2014. Mixed infection by Legionella pneumophila in outbreak patients. Int J Med Microbiol 304:307–313.

32. Diederen BM, de Jong CM, Aarts I, Peeters MF, van der Zee A. 2007. Molecular evidence for the ubiquitous presence of *Legionella species* in Dutch tap water installations. J Water Health 5:375–383.

33. Rivera JM, Aguilar L, Granizo JJ, Vos-Arenilla A, Gimenez MJ, Aguiar JM, Prieto J. 2007. Isolation of *Legionella species*/serogroups from water cooling systems compared with potable water systems in Spanish healthcare facilities. J Hosp Infect 67:360–366.

34. Storey MV, Langmark J, Ashbolt NJ, Stenstrom TA. 2004. The fate of legionellae within distribution pipe biofilms: measurement of their persistence, inactivation and detachment. Water Sci Technol 49:269–275.

35. Gaia V, Fry NK, Afshar B, Luck PC, Meugnier H, Etienne J, Peduzzi R, Harrison TG. 2005. Consensus sequence-based scheme for epidemiological typing of clinical and environmental isolates of Legionella pneumophila. J Clin Microbiol 43:2047–2052.

36. Alikhan NF, Petty NK, Ben Zakour NL, Beatson SA. 2011. BLAST Ring Image Generator (BRIG): simple prokaryote genome comparisons. BMC Genomics 12:402.

37. D’Auria G, Jimenez-Hernandez N, Peris-Bondia F, Moya A, Latorre A. 2010. Legionella pneumophila pangenome reveals strain-specific virulence factors. BMC Genomics 11:181.

38. Price MN, Dehal PS, Arkin AP. 2010. FastTree 2-approximately maximum-likelihood trees for large alignments. PLoS One 5:e9490.

39. Cheng L, Connor TR, Siren J, Aanensen DM, Corander J. 2013. Hierarchical and spatially explicit clustering of DNA sequences with BAPS software. Mol Biol Evol 30:1224–1228.

40. Didelot X, Wilson DJ. 2015. ClonalFrameML: efficient inference of recombination in whole bacterial genomes. PLoS Comput Biol 11:e1004041.

41. Parks DH, Mankowski T, Zangooei S, Porter MS, Armanini DG, Baird DJ, Langille MG, Beiko RG. 2013. GenGIS 2: geospatial analysis of traditional and genetic biodiversity, with new gradient algorithms and an extensible plugin framework. PLoS One 8:e69885.

42. Jombart T, Ahmed I. 2011. adegenet 1.3-1: new tools for the analysis of genome-wide SNP data. Bioinformatics 27:3070–3071.

